# CineFinch: An animated female zebra finch for studying courtship interactions

**DOI:** 10.1101/2022.11.02.514933

**Authors:** Nikhil Phaniraj, Sanjana Joshi, Pradeepkumar Trimbake, Aditya Pujari, Samyuktha Ramadurai, Shikha Kalra, Nikhil Ratnaparkhi, Raghav Rajan

**Author notes:** These authors contributed equally (Aditya Pujari, Samyuktha Ramadurai, Shikha Kalra). Correspondence should be addressed to Raghav Rajan.

## Abstract

Dummies, videos and computer animations have been used extensively in animal behaviour to study simple social interactions. These methods allow complete control of one interacting animal, making it possible to test hypotheses about the significance and relevance of different elements of animal displays. Recent studies have demonstrated the potential of videos and interactive displays for studying more complex courtship interactions in the zebra finch, a well-studied songbird. Here, we extended these techniques by developing an animated female zebra finch and showed that ~40% of male zebra finches (n=5/12) sing to this animation. To study real-time social interactions, we developed two possible methods for closed loop control of animations; (1) an arduino based system to initiate videos/animations based on perch hops and (2) a video game engine based system to change animations. Overall, our results provide an important tool for understanding the dynamics of complex social interactions during courtship.

**SUMMARY STATEMENT:** We develop and test an animation of a female zebra finch to study song and courtship interactions in the male zebra finch.

## INTRODUCTION

Complex behavioural displays are used by animals to communicate with each other (Bradbury and Vehrencamp, 2011). These behavioural displays carry information necessary for successful communication and often, have multiple different components that are produced together. For example, the courtship dance of the blue-capped cordon bleu, a songbird, involves multiple rapid foot and head movements that occur just before song, a vocal signal (Ota et al., 2015). Are all of these different components (foot, head movements and vocal signal) important for successful courtship? Do individual components of the display carry information about courtship potential or is courtship potential signalled only by presence of the entire display? Answering these questions requires complete control over one of the interacting animals to ensure that different components can be produced independent of each other. Such control is provided by robotic dummies, videos and computer animations and animal behaviour studies have a rich tradition of using such stimuli (Tinbergen, 1948; Woo and Rieucau, 2011). For example, Patricelli and colleagues used robotic female satin bowerbirds in artificially staged courtship interactions to demonstrate that male satin bowerbirds regulate the intensity of their courtship displays based on female responses (Patricelli et al., 2006). Van Dyk and Evans used computer animations of Jacky dragon lizards to understand the dynamics of aggressive encounters and showed that lizards use multiple signals to assess the level of aggression (Van Dyk and Evans, 2008).

Recent studies have extended the use of dummies and videos to understand more complex interactions like courtship in songbirds. In the zebra finch, a well-studied songbird native to Australia, courtship involves a song and a dance by the male (Sossinka and Böhner, 1980; Ullrich et al., 2016; Zann, 1996). Females also respond with vocalizations and tail-quivering displays (Zann, 1996). Male zebra finches sing to taxidermically stuffed female zebra finches and to videos of female zebra finches and the characteristics of these songs are highly similar to courtship song directed at a live female bird (Bischof et al., 1981; Galoch and Bischof, 2007; James et al., 2019). In addition to courtship interactions, male zebra finches also interact with juvenile zebra finches during tutoring sessions. Juveniles learn songs more accurately from a live tutor than from song playbacks from a speaker (Derégnaucourt et al., 2013). This suggests that the visual stimulus of the tutor and possibly social interactions with a tutor are also important for accurate learning. While the mere presence of a visual tutor, in the form of a video, is not sufficient to enhance learning (Varkevisser et al., 2022a), a recent study showed that a robotic zebra finch that vocally interacts with juvenile birds does enhance learning (Araguas et al., 2022). Importantly, the robotic zebra finch provided closed-loop vocal interactions, i.e. vocal interactions were provided only when juveniles interacted with the robot and the timing of these vocal interactions were comparable to the timing of interactions with live tutors. The robotic zebra finch highlighted the potential of appropriately manipulatable, artificial stimuli for probing complex social interactions, but such stimuli have not been tested in the context of courtship.

In addition to robots, animations provide an attractive method to provide closed-loop social interactions. The advent of fast computer hardware and user-friendly open-source graphics software has made it much easier to generate and control animations (Stowers et al., 2017). They provide a complementary approach to robots with considerable flexibility for studying social interactions. Here, we developed an animation of a female zebra finch and showed that ~40% of male zebra finches tested, sang to this animation. We also demonstrated two possible ways to interactively control these animations.

## MATERIALS AND METHODS

All experimental procedures conducted were approved by IISER Pune’s Institute Animal Ethical Committee (IAEC) and were in accordance with the guidelines of the Committee for the Purpose of Control and Supervision of Experiments on Animals (CPSCEA), New Delhi. Zebra finches (n=25 males and n=5 females) were procured from a local vendor (n=7) or bred in our colony at IISER Pune (n=23). Birds bought from an outside vendor were used for experiments only after they had been in our colony at IISER Pune for more than 30 days. Birds were raised in individual cages (120 cm x 50 cm x 50 cm cage, up to 6 birds per cage) along with other birds of the same gender. Light conditions were regulated to maintain a 14/10 hour day/night cycle. *Ad libitum* access to food and water were provided at all times, unless otherwise mentioned. All birds were >100 dph at the time of the experiment (males > 100dph and females > 300dph).

### Experimental Apparatus

The apparatus consisted of a metal double-cage (46 cm x 23 cm x 23 cm) with each half separated by a glass slab (23 cm x 23 cm x 0.3 cm). One half housed the subject male bird. During live female trials, the other half housed a stimulus female bird. During video playback or animation trials, the other half had a Samsung galaxy tab S4 placed upright lengthwise with its screen touching the glass slab. To record song produced by the male, we placed a microphone (AKG Acoustics C417PP omnidirectional condenser microphone) on top of the cage above the male. A camera (GoPro Hero7) was placed outside the apparatus to record videos of the sessions.

### Experimental subjects

For the experiments where males were presented with videos of females (video trials), we used 10 male zebra finches for the 30s trials and 10 males for the 4-minute trials. For the animation experiments, we used 12 males. 4 males were common across the 30s and 4-minute trials, 1 male was common across both 4 minute trials and animation trials and 1 bird was common for 30s and 4-minute video trials and animation trials. For the video experiments, we used 3 female zebra finches for the 30s trials and 2 female zebra finches for the 4-minute trials. For the animation experiments, we used 2 females. One female was common across 4-minute and 30s video trials and one female was common across 30s video and animation trials.

### Stimulus videos

4 minute long videos (2560 x 1440 pixels @60 fps) of a female bird were recorded while the female interacted with a male bird (see Movie 1 for part of a video). The same experimental setup that was used for video trials with the male was used for this and the male was positioned just beside the video camera, on the other side of the glass slab (Fig. 1A). The aspect ratio of the video was fixed at 15.5 cm x 8.7 cm to ensure that the female bird in the video appeared life-sized.

**FIGURE 1.**
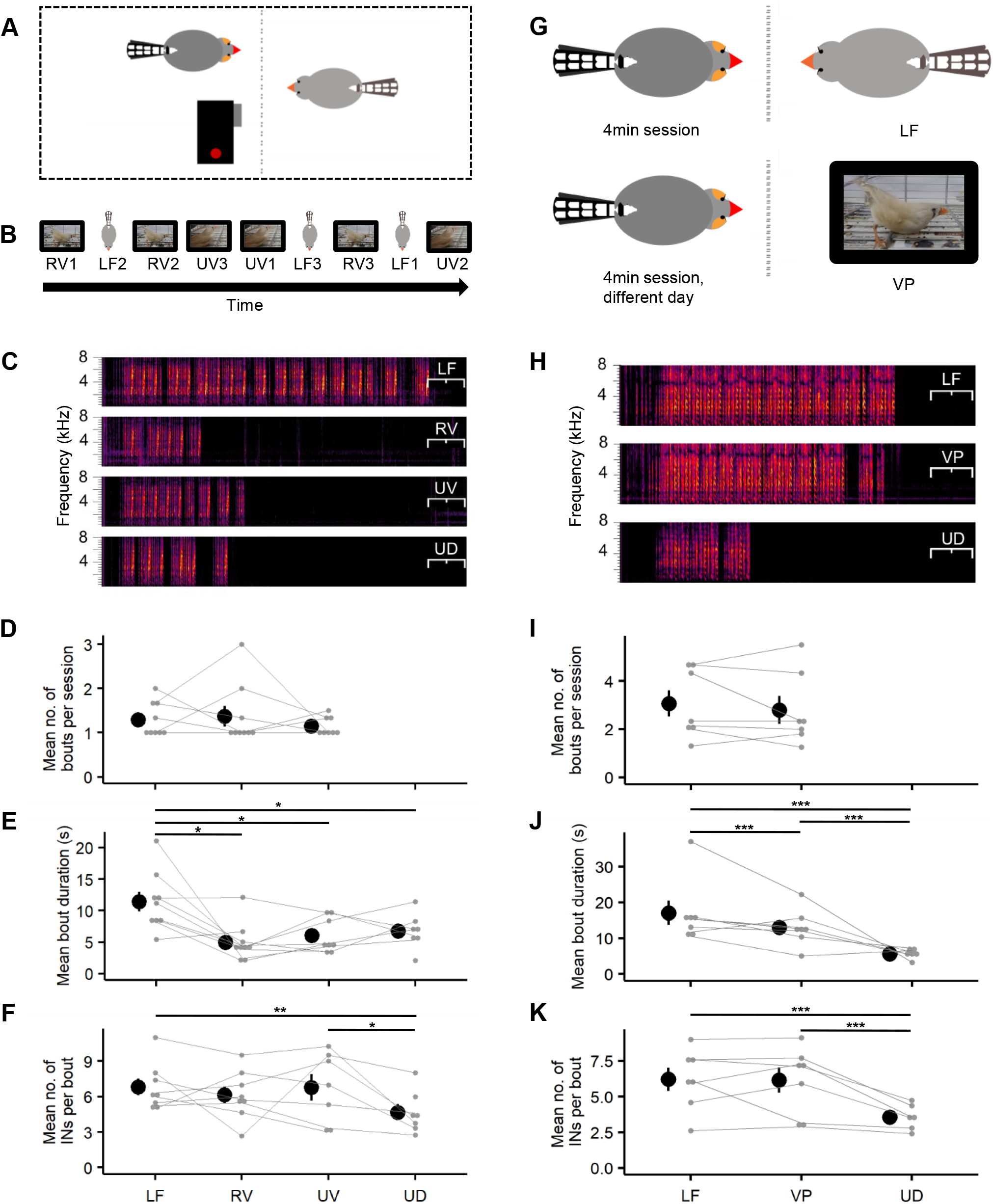
Male zebra finches sing longer song bouts to 4 minute videos as compared to 30s videos. A) Schematic depicting the way in which videos of females were recorded. The male is on the left side and the female is on the right side, separated by a glass partition. A camera was placed near the male to record videos of the female. (B) Order of 30s trials in a session for one representative bird. LF represents “live female”, RV represents “responsive” video and UV represents “unresponsive” video. Inter-trial-time of 5 minutes was used. (C) Example spectrograms showing songs produced by one male for the 3 different kinds of stimuli and undirected singing (UD) when the bird was alone. (D), (E), (F) mean number of song bouts per session (D), mean song bout duration (E) and mean number of introductory notes (INs) (F) for all birds for the different trial types. Grey circles represent data for individual birds and grey lines connect data from the same bird. Black circles and whiskers represent mean and s.e.m across all birds (n=9 birds). (G) Schematic depicting the experimental protocol for long duration experimental trials (4 minute trials). LF represents live female and VP represents video presentation. (H) Example spectrograms showing the songs produced by one male for the 2 different kinds of stimuli and undirected singing (UD) when the bird was alone. (I), (J), (K) mean number of song bouts per session (I), mean song bout duration (J) and mean number of introductory notes (INs) (K) for all birds for the different trial types. Grey circles represent data for single birds and grey lines connect data from the same bird. Black circles and whiskers represent mean and s.e.m across all birds (n=7 birds). * represents p < 0.05, ** - p < 0.01, *** - p < 0.001 - Type II Wald Chi Square test followed by pairwise comparisons using estimated marginal means with Holm correction.

### Construction of Animations

Initially, a 13.67s long animation of a female zebra finch was made using Blender 2.90.1 (https://www.blender.org/). This initial animation consisted of 820 frames which were played at 60fps. To make this, first, a 3D model of a female zebra finch was sculpted (Fig. 2A). Next, colours from a photograph of a female zebra finch were extracted and used to paint the skin of the 3D model (Fig. 2B). A feathery texture was added. 2 core bones (grey in Fig. 2C) and 9 structural bones (yellow in Fig. 2C) were rigged into the body and 5 control bones (blue in Fig. 2C) were placed outside that connected to a single bone or multiple distant bones for synchronized body movements and provided for puppeteerlike control of the zebra finch (Fig. 2A-2D). One of the control bones was placed in front of the beak and the beak bone programmed to always point towards the beak control bone. The beak-head joint was kept rigid whereas the head bone was free to rotate around the neck bone giving the head-neck joint 3 degrees of freedom. The second control bone was placed behind the tail and the tail bone programmed to always point to the tail control bone. The tail bone could rotate around the joint in both horizontal and vertical plane but not around its long axis, giving it 2 degrees of freedom. 2 of the control bones were placed in front of the claw and the claw bones were programmed to always point to the claw control bone. 2 additional bones were placed behind the knee, outside the body, to mark endpoints for inverse kinematics of the knee (red in Fig. 2C). These two external bones prevented the knee joints from displacing beyond the marked endpoints. The ankle joint was kept rigid whereas the knee and the pelvis were allowed restricted rotation in only the vertical plane giving them 1 degree of freedom. The 5th control bone was the master control bone (deep blue in Fig. 2C) placed directly below the centre of mass of the bird and this bone connected the other 4 control bones and allowed for coordinated movements of the two legs, tail, and head.

**FIGURE 2.**
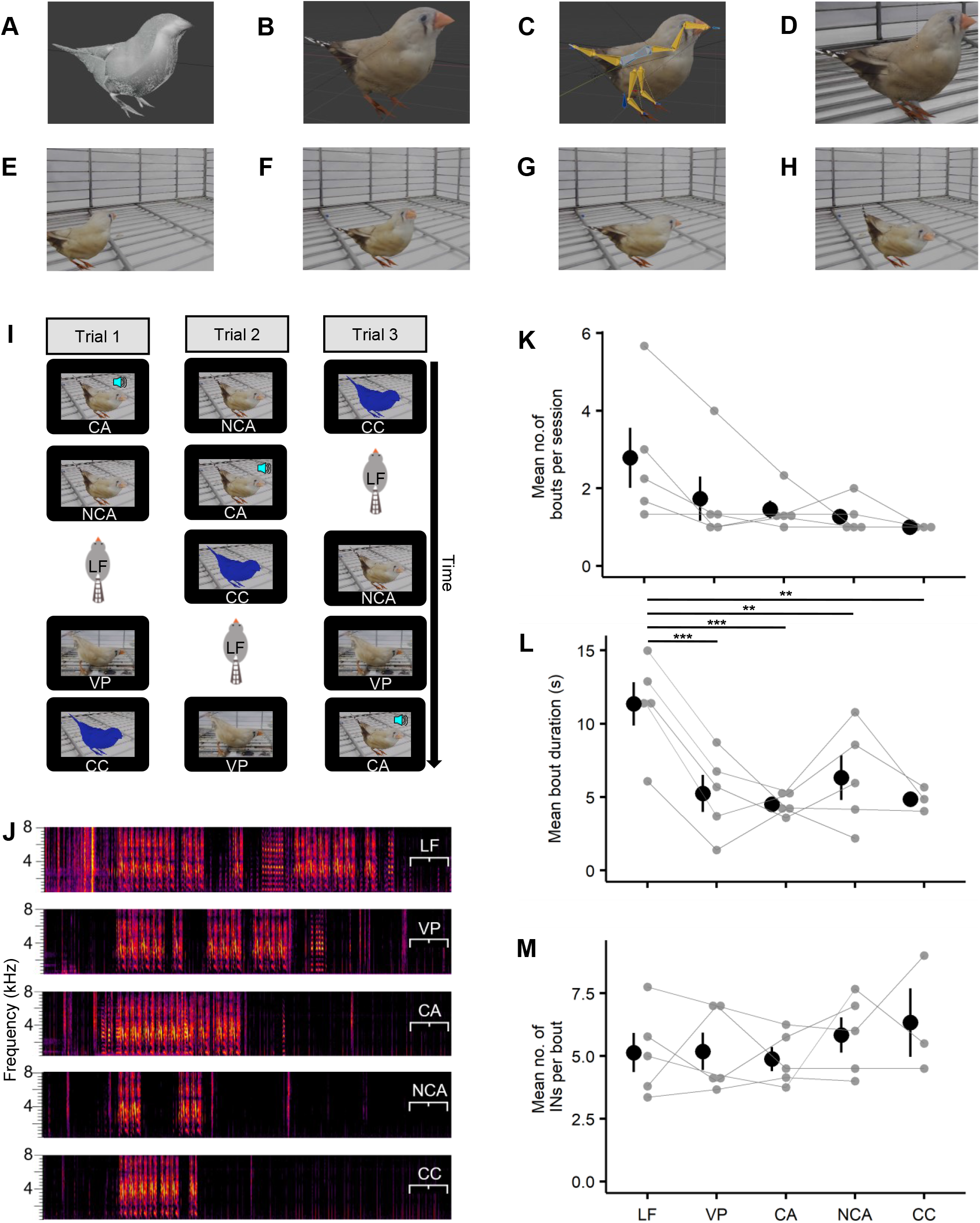
Construction and validation of an animated female zebra finch. (A), (B), (C), (D) Different stages in the construction of an animated female zebra finch in Blender; 3D model (A), with colour and feather texture (B). Positions of control bones with yellow representing structural bones, grey representing core bones, blue representing control bones, deep blue representing the master control bone and red representing the knee stop bones (C) (see Methods for details) and final image with empty cage (D). (E), (F), (G), (H) represent different animation frames with the female hopping into view, looking to either side and quivering its tail. I) Order of 2 minute experimental trials across 3 sessions for one representative bird. LF represents live female, CA represents call with animation, NCA represents no call animation, VP represents video presentation and CC represents colour control animation. Inter-trial-time of 5 minutes was used. (J) Example spectrograms showing the songs produced by one male for the 5 different kinds of stimuli. (K), (L), (M) mean number of song bouts per session (K), mean song bout duration (L) and mean number of introductory notes (INs) (M) are shown for all birds for the different trial types. Grey circles represent data for individual birds and grey lines connect data from the same bird. Black circles and whiskers represent mean and s.e.m across all birds (n=5 birds). * represents p < 0.05, ** - p < 0.01, *** - p < 0.001 - Type II Wald Chi Square test followed by pairwise comparisons using estimated marginal means with Holm correction.

The initial animation started with the female zebra finch hopping in from the left, performing 2 head turns (towards its right and its left), hopping to the centre, rotating towards the viewer, bending, and tail quivering (see Fig. 2E-2H for frames from the animation). After the tail quivering, the female was made to go back to the initial posture, hop to the right, perform 2 head turns (towards its right and its left), and hop to the right again – out of view. The velocity of hops, head turns and tail quivering were roughly similar to those of the female in one of the 30s responsive videos. We only matched the velocity qualitatively and did not reproduce exact statistics of movements. This initial animation was duplicated, laterally inverted and stitched to the end of the original animation to make the female enter from the left, exit from the right, then enter from the right, and exit from the left. This entire animation sequence, now 1640 frames (27.33s) long was again duplicated thrice and the 4 parts stitched together. A 10s clip showing an image of an empty cage was added to the beginning and the entire animation video was ~ 2mins (119.33s) long. A 5s long audio clip with 3 female calls was added to the first 5s of the animation (see Movie 2). The final animation video was played with the audio on (Call animation) or muted (No call animation). A color control animation was made where the bird in the animation was entirely painted with deep blue and the feathery texture removed (see Movie 2). Everything else was kept the same.

### General experimental procedures

All birds used for experiments were housed singly in small cages (23 cm x 23 cm x 23 cm) starting a few days before the start of the experiment and for the entire duration of the experiment, unless otherwise mentioned. For 30s trials, pairs of male subject birds were acclimatized to the setup by housing them in the apparatus – one bird on each half of the setup for 48 hours – 2 weeks before the start of the experimental trials, and again for 48 hours, before the time of the experimental trial. All experimental trials were conducted in sound-attenuation enclosures (Newtech Engineering Systems, Bengaluru or custom-made enclosures). For all experiments, songs were recorded with a microphone placed at the top of the cage. Signals from the microphone were amplified using a mixer (Behringer XENYX 802) and then digitized on a computer system at a sampling rate of 44,100 Hz using a custom-written Python based software. Undirected songs were recorded over a period of 24 hours, with birds housed singly in a cage (23 cm x 23 cm x 23 cm) in the same sound isolation box. 1 of the 7 birds never sang undirected songs.

### Video presentation procedures

Across all trial types, videos were played manually by the experimenter, without the audio, as a window on the screen. The video window overlapped an image of an empty cage in the background. The tablet was placed in its position as soon as the video was started. Once the video ended, the video window disappeared showing only the image of the empty cage on the screen. Animations were also converted into videos as described above and played using the same procedure.

### Short duration experimental trials (30s Video trials)

For the 30s video trials, experimental trials were conducted over a period of 10 days with one bird going through the experiment each day. Two, 30s long, silent, videos of each female were chosen as stimuli; (1) a “responsive” video where the female showed tail quivering and (2) an “unresponsive” video where the female did not show tail quivering. These videos were cut out of the 4 minute long videos of female birds that were recorded as described above. Along with the “responsive” and “unresponsive” videos, we also presented a live female as a third stimulus. Each of the three stimuli was repeated thrice and the 9 stimuli were presented to the male in random order separated by 5 minutes (Fig. 1B).

### Long duration experimental trials (4-minute video trials)

For the 4-minute video trials, we used the 4 minute long videos recorded as described above. Experimental trials were conducted over multiple days. On a given day, male birds were briefly checked with ~10s exposure to the tablet to see if they sang to the video. Birds went through the trial only if they sang to the video during the brief exposure (n=3/10 birds did not sing to the video on their first exposure and were not used further). During the actual trial, the subject bird was exposed to either the live female or the video for 4 minutes. For the live female condition, one of the two females was randomly chosen as the stimulus. For the video playback condition, one of the two videos (one of each female stimulus) was chosen to be played. For each male, only one type of trial was conducted on a given day. This procedure was repeated for each male on separate days until we obtained atleast 6 song bouts for each trial type (median of 7 days per male; range: 6-18).

### Animation trials

For the animation trials, birds were tested with a single exposure to the 2 minute animation of the female zebra finch without audio. Only 5/12 birds tested sang to this animation and these 5 birds were chosen for further experiments. Experimental trials consisted of 3 days of experiments with each day consisting of 5 types of trials (live female, video of female, animation with call, animation without call and colour control animation) in a random sequence. Each trial lasted 2 minutes and trials were separated by 5 minutes.

### Song Analysis

Songs were analysed as described earlier (Kalra et al., 2021; Rajan and Doupe, 2013; Rao et al., 2019), using custom-written scripts on MATLAB (MathWorks). Briefly, songs were first segmented into syllables based on an amplitude threshold. Syllables with inter-syllable intervals < 5 ms were merged into one syllable and syllables shorter than 10s were discarded. Then, syllables were manually labelled based on their appearance on a spectrogram; similar looking syllables were given the same label. The sequence of syllables that repeated in a bird’s song was identified as a motif (Sossinka and Böhner, 1980; Zann, 1996). Short syllables that were repeated a variable number of times at the beginning of a bout were considered as introductory notes (Kalra et al., 2021; Rajan, 2018; Rajan and Doupe, 2013; Rao et al., 2019). The number of introductory notes at the beginning of a bout was quantified by starting with the introductory note immediately preceding the first syllable of the bout and counting backwards until either (1) the previous note was not an introductory note or (2) there was a silence > 500 ms (Kalra et al., 2021; Kao et al., 2005; Rajan and Doupe, 2013; Rao et al., 2019; Sossinka and Böhner, 1980). Song bouts were defined as groups of vocalizations that contained at least one motif syllable and were separated from other such groups by more than 2 s of silence. All song bouts sung by birds towards live females and video/animation were considered for analysis as independent song bouts and were combined with all other song bouts across sessions/days. Of all the undirected bouts sung by a bird, 15 were randomly chosen for analysis for the 4 minute trials (median: 15; range: 14-16) and 4 were chosen randomly for analysis for the 30s trials (median: 4; range: 2-6). The number of undirected bouts to analyse was chosen to be comparable to the number of song bouts produced by the male in response to the live female and video trials.

### Tutoring of birds with videos

Juveniles used for tutoring were isolated as described earlier (Kalra et al., 2021). Briefly, fathers were removed from the nest around 12 days post hatching (range: 9-13) and juveniles were housed along with their mother until they could feed on their own (range: ~35-40 days post hatch). Following this, individual juveniles were housed singly and tutoring began on ~37dph (range: ~ 36-46 dph). Tutoring was done in a cage that contained two perches; one was active and associated with playback of the tutor song as described below and one perch was a dummy. The active perch had an IR beam running across it, which was monitored by an Arduino board (https://www.arduino.cc/). When the bird jumped onto the perch a break in the IR beam was detected by the Arduino-MATLAB interface. This triggered the tablet screen to play the video of the male bird along with audio playback of the tutor song. Specifically, an IR-beam break initiated a command to virtually “tap” on a specific coordinate on the tablet screen. This “tap” opened the video in a VLC window of whose size was chosen to ensure that the bird in the video was of a realistic size. The latency of the perch-hop to display loop was generally <1 sec but we noticed that it was longer for the first few hops in a session. After the video ended, the display was reset to an image of the far end of the cage as it would have appeared if the tablet were not present. This was also used as the initial image before tutoring began. Tutoring was done for 1-2 sessions per day (median number of tutoring sessions across birds = 45; range = 24-51; some days, we did not conduct tutoring). In each session, we limited the number of possible song playbacks to 20 as too much exposure to song has been shown to result in poor imitation (Tchernichovski et al., 1999). Each session lasted until the bird had exhausted the quota of 20 playbacks. If the bird did not trigger 20 playbacks, then the session was terminated after 1 hour. Passive playbacks were used occassionally to prompt the bird to hop on the perch. Playback songs consisted of introductory notes (INs) followed by 2 motifs (n=3 were tutored with 2 INs and n=3 were tutored with 7 INs). At ~70 dph (range: 63-79 dph), we switched the number of INs for the tutor song and tutoring with the changed song continued till 90 dph. These were done to determine if the number of INs can be changed during the course of learning and are part of another ongoing study. However, for the purpose of this study, we only considered the accuracy of imitation of the motif and this motif was not changed during the entire duration of tutoring.

Song similarity was calculated using 5 example motifs recorded from these birds at ~90 dph (range: 90-100 dph). Similarity calculations were done using Sound Analysis Pro (SAP) (Tchernichovski et al., 2000) as described earlier (Kalra et al., 2021). Briefly, asymmetric time-course similarity measurements were made between tutor song and 5 example motifs and the average of these similarity values was taken as a measure of similarity to the tutor song. For comparison with birds tutored without videos, we used birds tutored at IISER Pune for a different study (n=10) (Kalra et al., 2021). These birds were tutored with 2 motifs without INs using active playback methods.

### Closed-loop animations

The animated female zebra finch described earlier was used in conjunction with a video game engine to create interactive animations. First, animations were added to the female model using keyframing, a technique used in blender. Animations included movements like hops, neck movements, body rotation, tail quivering, and panting. The animation also included female calls (3 calls per stimulus). The animated female was then exported in FBX (filmbox) format. FBX format was chosen for compatibility with the game development software, Unreal Game engine 4 (https://www.unrealengine.com/en-US). This model was imported into Unreal engine 4 (version 4.27.2). Unreal engine 4 was used to create a simple game for generating potential stimuli. First, we created a 1D blend space, i.e. an axis that allowed us to gradually blend a stationary animated female zebra finch into a hopping, tail-quivering zebra finch. Then, key-presses were associated with different positions along this axis, thereby allowing us to control the speed of hopping and tail-quivering. The key-press associations were made within the code of the game engine (Fig. S3) and such stimuli generated from the game would allow interactive-control of the animation in real-time.

### Statistical analysis

We did not perform any apriori sample size calculations. All statistical analyses was done with R. Sample sizes used were comparable to other studies (Ikebuchi and Okanoya, 1999; James et al., 2019; Takahasi et al., 2005) Generalized Linear Mixed Models (GLMMs) were used to statistically analyse the data. We considered each song bout as a single unit and considered all song bouts across trial types for statistical analyses. Response variables were fit using the appropriate distributions; bouts per presentation and number of INs were count variables and were fit using a Poisson distribution, bout durations were skewed towards lower duration values, so we log transformed these values and a log-linked Gaussian distribution was used. Bird identity was a random effect and stimulus type was a fixed effect for all analyses. Day Number, Presentation number and bout number (all ordinals) were included as fixed terms while analyzing the number of INs and bout durations in the animation trials. Only day number and presentation number were included as fixed terms (along with stimulus type) for analyzing the number of bouts per stimulus for animation trials. Only presentation number and stimulus type were fixed terms for analyzing the number of bouts per stimulus for the 30 s video trials. For the rest, stimulus type was the only fixed term (day, presentation, and bout numbers could not be included when comparisons with undirected songs were made). Type II Wald Chi Sq. tests were used to determine statistical significance followed by pairwise comparisons between stimulus type using estimated marginal means with Holm correction. To compare the log ratio of time spent singing across different conditions, and to analyze the number of bouts per stimulus in the 4-minute video trials, we used a Kruskal-Wallis test. An alpha level of 0.05 was considered for all statistical analyses.

## RESULTS

### Male zebra finches sang short song bouts to 30s videos of female zebra finches

Our goal was to make an animated female zebra finch that could be used for understanding social interactions. We first tested our display systems by examining responses of male zebra finches to videos of female zebra finches. We used two different videos (see Fig. 1A and Methods for video recording procedures) of the same female: (1) a “responsive” video where the female made tail-quivering movements and (2) an “unresponsive” video without tail-quivering movements (Movie 1). Individual males were presented with 3 presentations each of the two types of videos and 3 presentations of a live female in random order (Fig. 1B; see Methods for details). Male birds sang comparable number of songs to videos of female birds and to live females (Fig. 1C, 1D, Movie 1, Fig. S1A, Fig. S2A). As reported earlier (James et al., 2019), song bouts directed to either of the videos were considerably shorter than song bouts directed to a live female (Fig. 1E). Songs directed towards live females and songs directed towards videos of females were preceded by greater number of introductory notes as compared to “undirected” songs produced when the bird was alone (Fig. 1F), indicating the “directed” nature of songs sung to the videos (Sossinka and Böhner, 1980). Overall, these results showed that males sing “directed” songs to videos of females and singing behaviour was not influenced by responsiveness of the female.

### Longer duration videos elicited longer song bouts

To test whether longer stimulus durations elicit longer song bouts, we next presented males with 4 minute videos consisting of both “responsive” and “unresponsive” segments. Each day, individual males were presented with either a video or a live female each day (Fig. 1G) and trials were conducted for multiple days in a random order until we obtained atleast 6 song bouts in both conditions (see Methods for details). Birds sang comparable number of song bouts to the live female and to the videos (Fig. 1H, 1I, S1C). Song bouts to the live female were still longer (Fig. 1J), but the difference in length of song bouts to videos and live females was smaller with longer video durations (Fig. S1C, S2C). This was largely a result of more singing after the first 30s (Fig. S1C). Songs directed to the 4 minute videos were also preceded by higher number of introductory notes compared to undirected songs (Fig. 1K). Overall, these results suggest that video duration plays a role in determining song bout duration possibly by giving the bird more time to detect and respond to the presence of a female on the screen.

### Building an animation of a female zebra finch

To obtain greater control of the behaviour, we next constructed an animation of a female zebra finch. The animation was constructed in Blender by sculpting a 3D model of a female zebra finch, painting the skin with realistic colours and adding a feather texture (Fig. 2A, 2B). Bones were rigged into the body to allow for puppeteer-like control of movement of specific body parts (Fig. 2C; see Methods for details). Finally, the bird was placed in a background of a cage (Fig. 2D). We used this to construct a simple animation that involved the female hopping in, performing a few head turns and quivering its tail. This sequence was repeated multiple times to construct a 2 minute video that began with the image of an empty cage (Fig. 2E – 2H; first 2 minutes of Movie 2; see Methods for full details). This video was played with the 5s of female calls on appearance of the female (Call animation) or without audio (No call animation). As a control, we used a colour-modified version where the entire bird was painted with deep blue and the feathery texture removed (last 2 minutes of Movie 2). Everything else including all the movements were kept the same.

### Male zebra finches produced courtship songs directed at the animations

To test the ability of this animation to elicit song, we tested male birds across 3 sessions (1 session/day) using 5 stimuli presented in random order in each session. The five stimuli used were (1) a live female, (2) a video of a female, (3) video of the animated female zebra finch with and without audio (3 and 4) and the (5) colour-modified control animation without audio (Fig. 2I; see Methods for details). Stimulus duration was kept at 2 minutes. We selected birds based on response to a single 2 minute exposure to the animation without calls. 5/12 males sang to this single exposure and were chosen for the 3 sessions. Male birds sang to all stimuli with comparable numbers of song bouts (Fig. 2J, 2K, S1B, S2B). Song bouts directed at the animation (Movie 3) were comparable in duration to song bouts directed at videos (Fig. S2D), but significantly shorter than song bouts directed at live females (Fig. 2L). Songs directed at animations began with a large number of introductory notes confirming the “directed” nature of these songs (Fig. 2M). Interestingly, 3/5 birds also sang to the colour-modified control animations (Fig. 2J, 2K, 2L). These bouts were also preceded by a large number of introductory notes and were comparable in length to bouts directed towards the animation (Fig. 2M). These results demonstrate the potential of using animations to elicit song and suggest that such animations could be used to further study courtship interactions in the zebra finch.

### Methods for closed loop control of the animation

To use these animations to study courtship interactions, we needed methods to change these animations in real-time, i.e. we needed a closed-loop system. Here, we used two different approaches to produce interactive animations; (1) using an Arduino to control the appearance of a video and (2) using the animation in conjunction with a gaming engine.

In the first approach, we used an Arduino (a single-board microcontroller) based approach to control the appearance of a video for tutoring young birds. Since young birds learn better from a live tutor than from song playback from a speaker (Derégnaucourt et al., 2013), we used active tutoring methods to test whether including a video of a male bird along with song playback increased copying accuracy. Juveniles could hop on a perch to elicit song playback and video playback of a male bird (Fig. 3A; see Methods for details of procedure). Juveniles tutored by this method (n=6) copied tutor songs with accuracies similar to juveniles tutored by song playback alone (Fig. 3B shows spectrograms of tutor song and songs of two juveniles; Fig. 3C; n=6 birds with video and n=10 birds with only audio playbacks from Kalra et al. (Kalra et al., 2021)). These results suggested that the visual presence of an adult bird, albeit on a screen, does not improve copying accuracy and is supported by similar results from a recent study (Varkevisser et al., 2022a).

**FIGURE 3.**
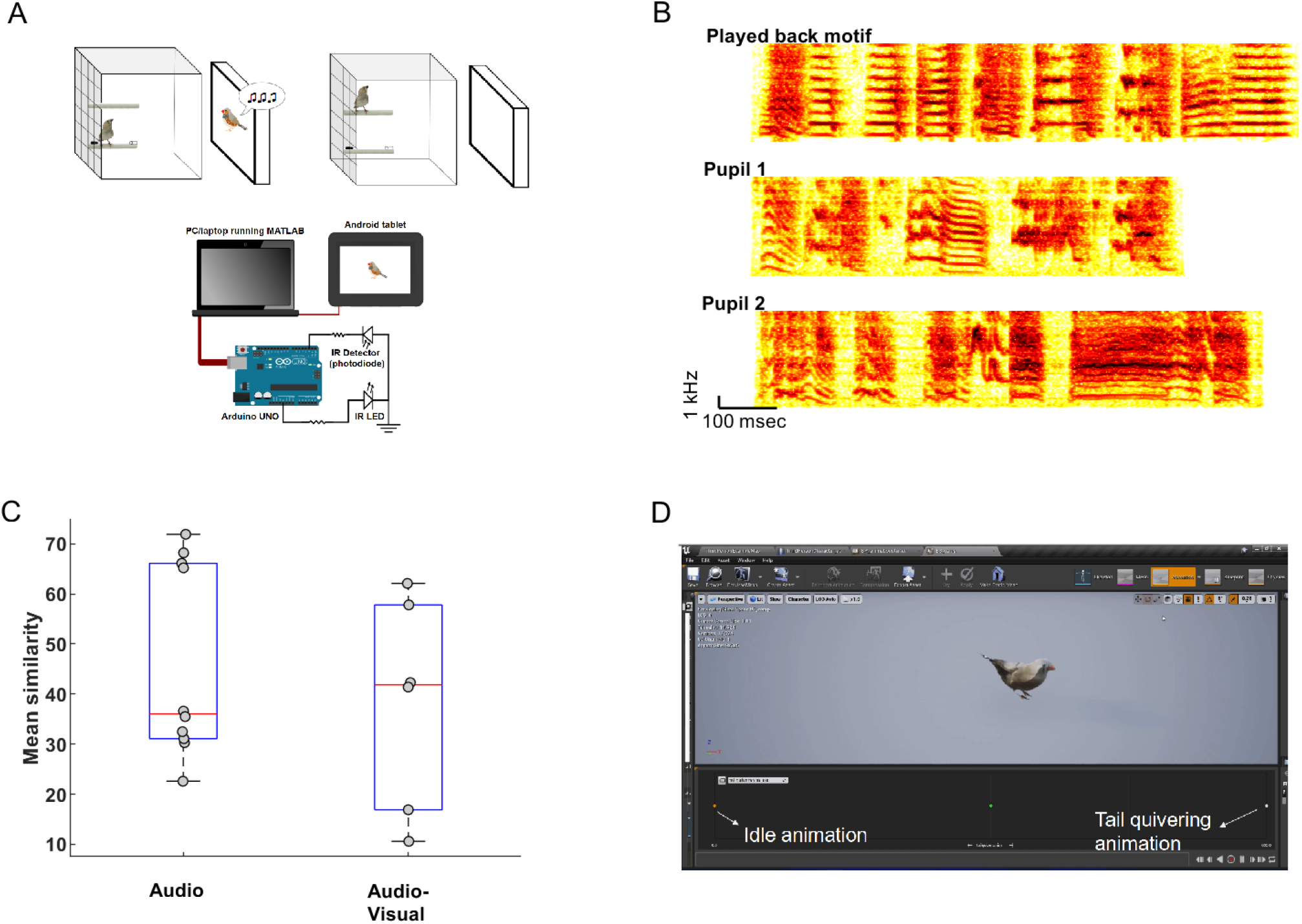
Potential methods for interactive control of the animation. (A) Schematic of experimental setup and Arduino based approach to control song audio and video playback based on perch hopping behaviour of a juvenile zebra finch. Briefly, hopping on a perch triggers playback of song audio and video through an Arduino-MATLAB connection. (B) Example spectrograms of tutor song and the songs of two juveniles tutored using this paradigm. (C) Boxplots showing the similarity to tutor song for juveniles tutored using only audio playbacks (subset of birds from an earlier study (Kalra et al., 2021)) and audio-visual playbacks. Circles represent data from individual birds. (D) A view of the 1D blendspace in Unreal Engine 4 with the axis extremes representing “An idle animation” and “Tail quivering animation”. Different positions along the axis represent different speeds of the tail quivering.

While, the Arduino based approach was simple and easy to use, real-time control of the animation was not possible, as latencies from perch hop to video playback were long (mean +/- std. was 0.74s +/- 0.16s). In our second approach, we used a video game engine, Unreal Engine 4 to construct interactive animations. We imported the female zebra finch from Blender into Unreal Engine 4 and created a 1D blend space, i.e. an axis that allowed us to gradually blend a stationary animated female zebra finch into a tail-quivering zebra finch (Fig. 3D, S3; see Methods for details). Different speeds of movement along the axis were connected to different durations of a “key press” on a regular keyboard. While, we connected “key-presses” to different states of the animation, it should be possible to use input from a microphone to control movement of the animation allowing for song-contingent control. Such methods have been used to make birds change the fundamental frequency of their vocalizations (Tumer and Brainard, 2007). Overall, these two methods provide the potential for generating closed-loop, interactive, animations for studying courtship dynamics.

## DISCUSSION

Here, we demonstrated potential use of animations to study courtship interactions in a songbird, the zebra finch. Specifically, we first showed that male zebra finches sang to videos of female zebra finches and song bouts were longer in response to longer videos. Second, we built an animation of a female zebra finch and showed that 42% of males sang to the animation. Finally, we showed the potential for real-time control with tools for interactive control of the animation; a feature that is essential for analysing courtship interactions in real-time. Overall, our results highlight the power of animations for studying courtship interactions in songbirds and provide a new tool for the same.

### Duration of the video is a factor that influences the amount of song produced

Artificial stimuli like animations, need to capture important details of the natural stimulus to be effective. What aspects of a video/animation stimulus are important for eliciting song from a male zebra finch? Our results show that stimulus duration is an important factor to consider. Male birds sang longer song bouts when presented with 4 minute videos as compared to 30s videos of females (Fig. 1). Both in earlier studies (James et al., 2019) and our experiments, videos were played without sounds. Live females call and so, when presented with a live female, male birds would experience both visual and auditory stimuli. Such multimodality is known to be essential for learning from a tutor (Varkevisser et al., 2022b) and multimodal signals are common in the courtship displays of other songbird species (Mitoyen et al., 2019; Ota et al., 2015). Given the absence of auditory cues in the videos, longer duration videos may give birds more time to notice the video and decide to sing. Integration of auditory cues may increase responses to videos. In line with this, a recent study also used audio and visual cues to successfully allow two birds to interact with each other virtually (Larsen et al., 2022).

### Improving responses to animations

Our animation successfully elicited songs from only 42% of the birds tested. What can be done to make more birds respond to animations? One possibility is to make animations more realistic by incorporating realistic movement parameters by using pose-estimation software like DeepLabCut to track, quantify movements of live females (Lauer et al., 2022; Mathis et al., 2018). Another possibility is to use colour corrections based on the bird’s visual system. Such colour-corrected videos have been shown to increase the attention of juvenile birds to videos of tutors (Varkevisser et al., 2022b). In our own experiments, the response of some birds to colour control animations suggests the possibility that the colour scale was not realistic enough.

### Outlook for using animations for closed-loop studies of social interactions

Our animations and the use of these animations in conjunction with video game engines (like Unreal Engine) provide a novel way to study social interactions in birds. Recent studies have used a robotic zebra finch for studying social interactions between juvenile birds and a tutor (the robotic bird) (Araguas et al., 2022). While a robot provides physical interactions that animations cannot provide, animations can be built and manipulated more easily. Animations, like the one we have developed would provide a complementary approach to robotic zebra finches to study social interactions using virtual-reality setups. A recent study used a virtual reality arrangement to show that two zebra finches can successfully interact with each other virtually (Larsen et al., 2022). Our animations and video game engines, combined with this setup could facilitate testing and analyses of social interactions in much greater detail.

### Broader applications of animated animals

Here, we focused on developing and controlling an animation for studying courtship interactions in zebra finches. Similar animations with other animal species can be useful for a diverse range of socio-cognitive and behavioural experiments. For instance, such animations could be used in social learning experiments to study the process of social learning. Such experiments typically require one animal in a group to learn a novel task asocially, following which social transmission of this learning can be studied (Heyes, 1994; Rendell et al., 2010; Van Schaik and Burkart, 2011). However, such asocial learning can take time. This can be speeded up using videos to teach the first animal a novel task. Animations could also be used to study interspecies encounters and animals’ responses towards heterospecifics (Gröning and Hochkirch, 2008; Peiman and Robinson, 2010; Sridhar and Guttal, 2018). In some species, response to heterospecifics is plastic and is observed to vary with heterospecific individuals (Lehtonen et al., 2010; Magellan, 2020). However whether physical features (size, body shape, etc.) or behavioural features (aggression, submission, etc.) are used by animals to identify heterospecifics remains unclear. Animations, such as ours, offer independent control over physical and behavioural features and would understand interspecies interactions. Overall, our methods will not only aid the study of animal courtship interactions but will also be extendable to other fields from animal cognition to ecology.

## SUPPLEMENTARY INFORMATION

Supplementary information includes 3 supplemental figures and 3 movies.

## DATA ACCESSIBILITY

All data, videos, animations and files related to animation are available on reasonable request from the corresponding author (RR -).

## ACKNOWLEDGMENTS

We would like to thank Prakash Raut for bird colony maintenance and Vrushali Rao Gumnur and Aboli Ektare for preliminary tests with different monitors and video stimuli. We would like to thank Hamish Mehaffey, Mimi Kao and Girish Deshpande for helpful comments on the manuscript and the Rajan lab for useful discussion related to the project.

## AUTHOR CONTRIBUTIONS

NP and RR designed the study. NP did the experiments with videos and created the animated female zebra finch in Blender. SJ carried out the animation related experiments. NR did some initial work on interactive animations using Unity and PT used Unreal engine 4 to construct interactive animations. SK, AP and SR constructed the tutoring setup and tutored juveniles with video and audio playbacks. NP and RR wrote the manuscript in consultation with SJ, PT, SK, AP, SR and NR.

## SUPPLEMENTARY INFORMATION

Supplementary information consists of 3 supplementary figures and 3 movies.

### SUPPLEMENTARY FIGURE LEGENDS

**FIGURE S1.**
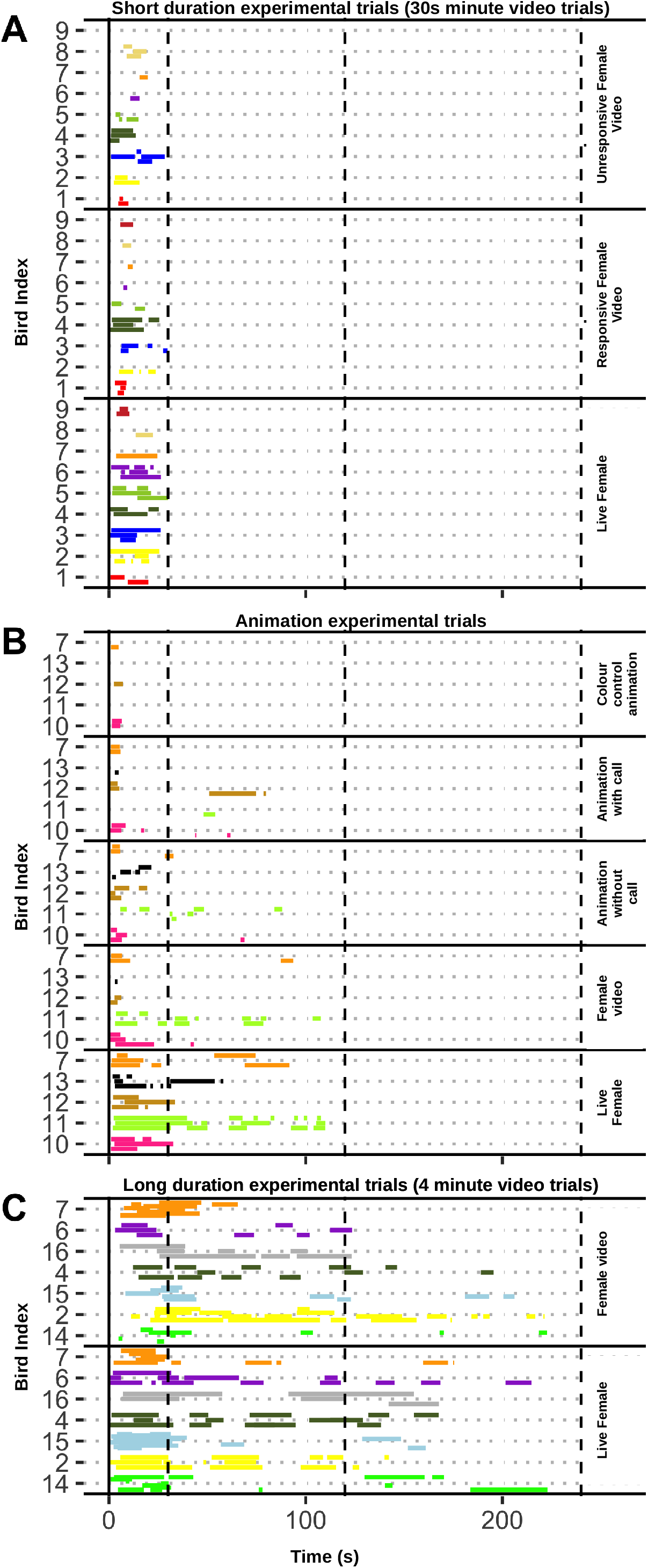
Raster plots showing song bouts produced by all birds during short-duration, long-duration and animation experimental trials. (A), (B), (C) Raster plots showing song bouts produced by all birds during short-duration (A), animation (B) and long-duration experimental trials (C). Each bar represents a song bout and the length of the bar represents the duration of the song bout. Different colours represent different birds. Bird indices are indices given to unique birds. Birds that were common across different experimental trials can be tracked using this bird index (for eg. bird #7 was used for all types of experimental trials).

**FIGURE S2.**
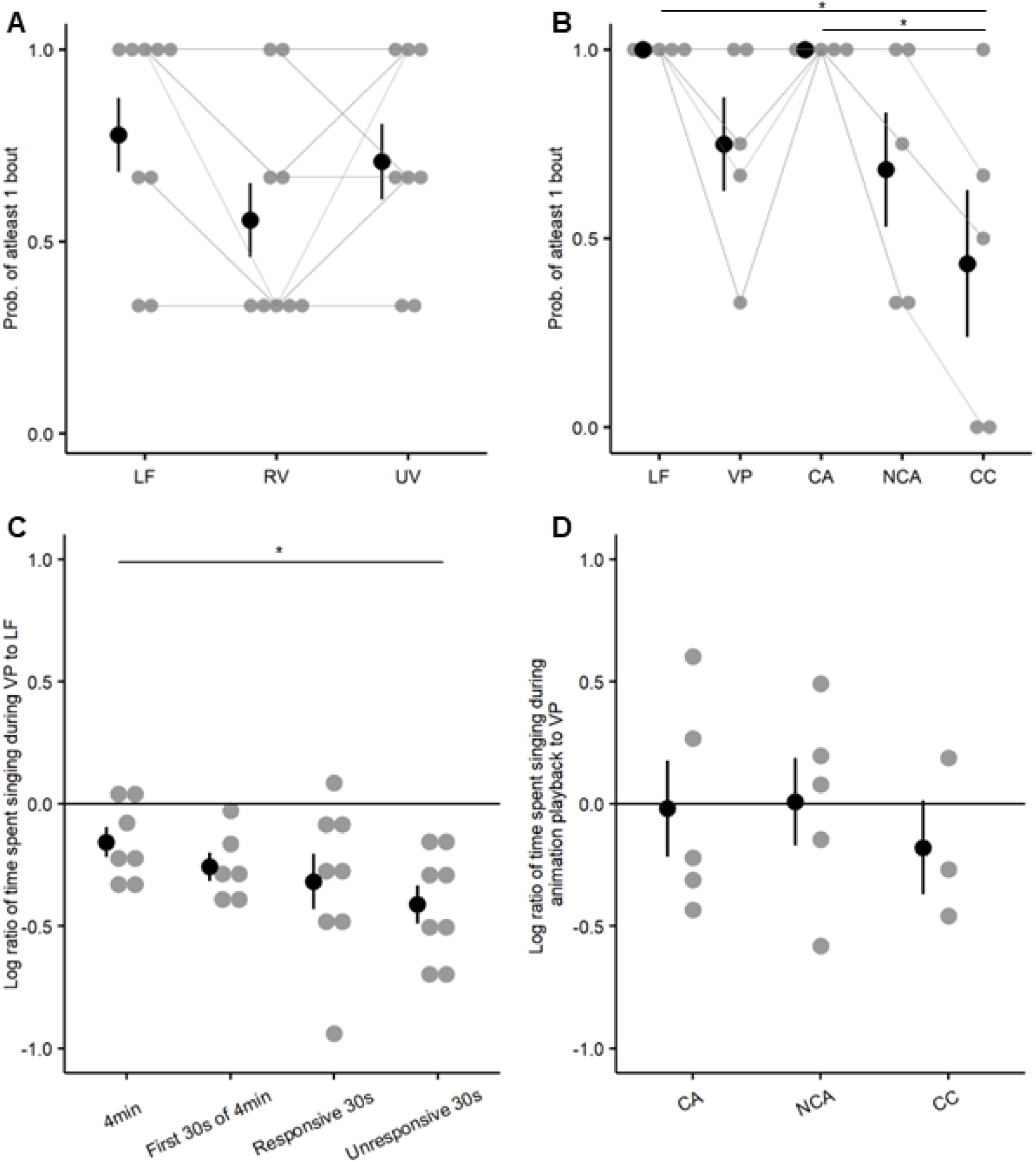
Probability and amount of song produced towards different stimulus types. (A), (B) Probability of singing atleast 1 bout of song for 30s video trials (A) and animation trials (B). (C) Log ratio of time spent singing to video presentations relative to live females. (D) Log ratio of time spent singing to animation relative to video presentation. Grey circles represent data from individual birds and grey lines connect data from the same bird. Black circles and whiskers represent mean and s.e.m across all birds.

**FIGURE S3.**
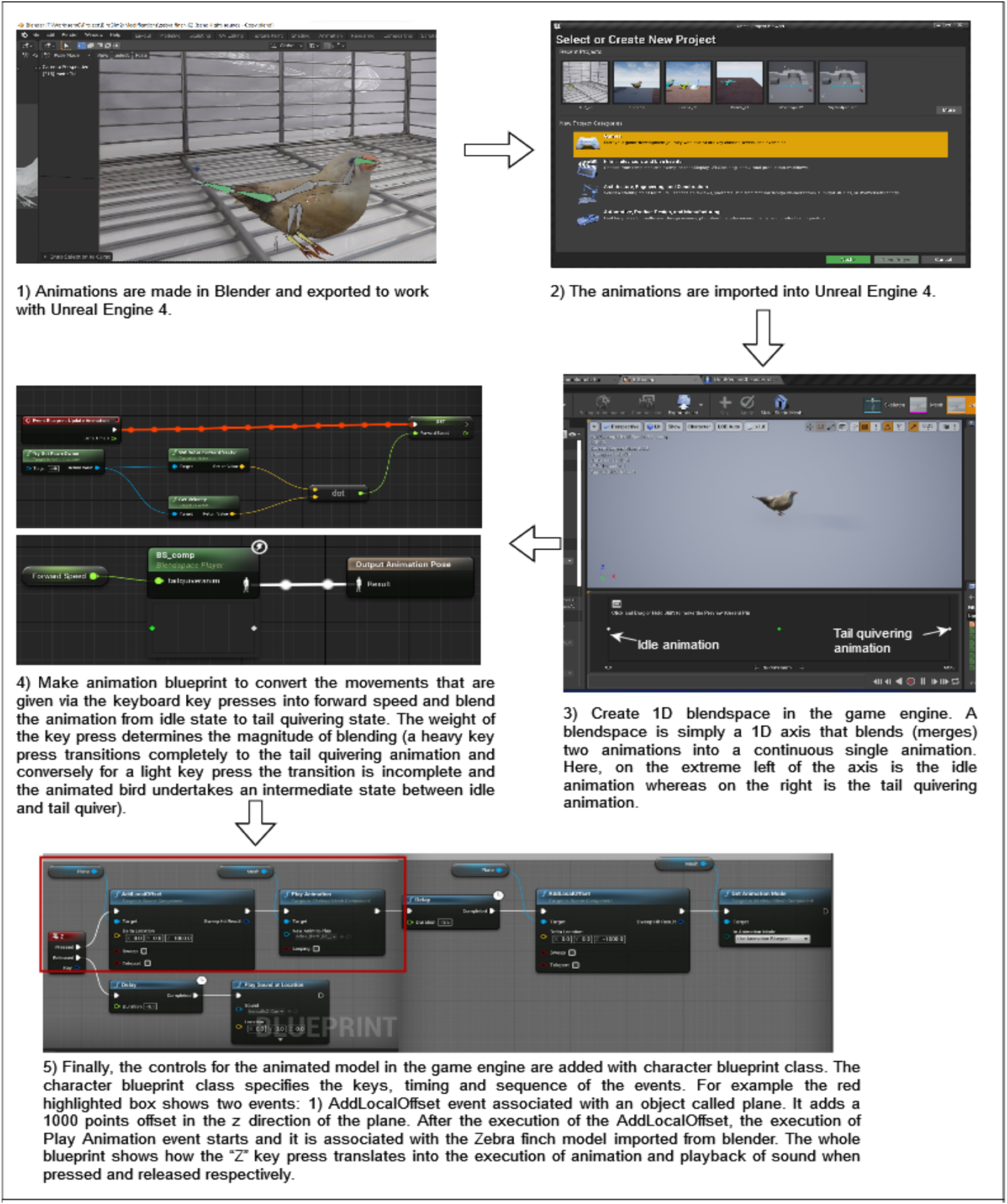
Construction of interactive animations using video game engine Unreal Engine 4. Flowchart depicting the process used to create interactive animations using the video game engine Unreal Engine 4.

### MOVIE CAPTIONS

**MOVIE 1 Examples of responsive and unresponsive videos and song bouts of a male zebra finch** Movie shows an example of a responsive video of a female (30s), an unresponsive video of a female (30s) followed by examples of a male singing to a live female and to a responsive video.

**MOVIE 2 Male zebra finches sing to animations of female zebra finches**

Movie shows one cycle of the animation and one cycle of the colour control animation. This is followed by examples of males singing to the animation and the colour control animation.

**MOVIE 3 1D blend space in Unreal Engine 4**

Movie shows a 1D blend space in Unreal Engine 4 with one end of the axis corresponding to an idle female zebra finch and the other end corresponding to a tail quivering zebra finch. Speed of tail quivering varies along the axis and an intermediate speed is also shown in the video.

